# An information-theory approach to geometry for animal groups

**DOI:** 10.1101/839548

**Authors:** Christoph D. Dahl, Elodie Ferrando, Klaus Zuberbühler

## Abstract

One of the hardest problems in studying animal behaviour is to quantify patterns of social interaction at the group level. Recent technological developments in global positioning system (GPS) devices have opened up new avenues for locating animals with unprecedented spatial and temporal resolution. Likewise, advances in computing power have enabled new levels of data analyses with complex mathematical models to address unresolved problems in animal behaviour, such as the nature of group geometry and the impact of group-level interactions on individuals. Here, we present an information theory-based tool for the analysis of group behaviour. We illustrate its affordances with GPS data collected from a freely interacting pack of 15 Siberian huskies (*Canis lupus familiaris*). We found that individual freedom in movement decisions was limited to about 4%, while a subject’s location could be predicted with 96% median accuracy by the locations of other group members, a pattern mediated by dominance, kin relations, sex, the relative time of the day and external events, such as feeding. We conclude that information theory-based approaches, coupled with state-of-the-art bio-logging technology, provide a powerful tool for future studies of animal social interactions beyond the dyadic level.

## Introduction

The combination of naturalistic observations in the field and systematic experimental testing in captive settings has long been the gold standard in studies of animal behaviour and cognition (Kummer, 1984). Behaviour, in turn, is usually collected by human observers, who follow a range of established observation techniques (Altmann, 1974) and experimental designs (Zuberbühler & Wittig, 2011). Although this combination has provided unprecedented progress, it has a number of inherent and highly problematic flaws. First, behaviour coding relies on human categorisation, which is inherently prone to bias. A common way to address this is to carry out inter-observer reliability tests, which remedies the problem to some extent. Second, behaviour experiments are often in interaction with humans, which can introduce ‘Clever Hans’ effects and other forms of cuing (Pfungst, 1911). Third, human-led behaviour studies are usually restricted to reporting from single individuals interacting with its socio-ecological umwelt, which typically excludes analyses at larger levels, including how individuals contribute to group-level behaviour and decisions, or how they are influenced by them (Altmann, 1974; Snowdon, 1983). As a consequence, current theories of animal social behaviour are largely based on data collected during pair-wise interactions of animals (Branson, Robie, Bender, Perona, & Dickinson, 2009; Dankert, Wang, Hoopfer, Anderson, & Perona, 2009 De Chaumont et al., 2012; Langford et al., 2006).

More recently, there have been efforts to studying patterns of social interactions at the group level (Shemesh et al., 2013; Strandburg-Peshkin, Farine, Couzin, & Crofoot, 2015), but this is hardly possible without moving away from subjectively coding animal behaviour by human observers. Bio-logging is among the most promising tools for quantifying animal behaviour without subjective human coding (Dahl, Wyss, Zuberbühler, & Bachmann, 2018; Gerencsér, Vásárhelyi, Nagy, Vicsek, & Miklósi, 2013). For example, GPS-based inertial sensor technology provides high-resolution data of movement patterns, which can be further processed towards automated behavioural classification systems and descriptions of group-level interactions (Ákos, Beck, Nagy, Vicsek, & Kubinyi, 2014; Dahl et al., 2018; Nagy, Akos, Biro, & Vicsek, 2010; Strandburg-Peshkin et al., 2015), even in the absence of image-based tracking (Dell et al., 2014; Rasch, Shi, & Ji, 2016; Shemesh et al., 2013). Although these data are relatively easy to obtain, describing social interactions in groups is computationally challenging, requiring mathematical models of interaction that are complex (Bialek et al., 2012; Couzin, Krause, Franks, & Levin, 2005; Couzin, Krause, James, Ruxton, & Franks, 2002; Lukeman, Li, & Edelstein-Keshet, 2010; Vicsek, Czirók, Ben-Jacob, Cohen, & Shochet, 1995).

Here, we provide a means to study the social and environmental determinants of group formation in free-ranging, group-living animals. We used information theory (Shannon, 1948) to determine the degree to which individuals influence each other in choosing their location and how this is driven by social and environmental determinants. We test this tool on a pack of 15 dogs (*Canis lupus familiaris*; Siberian husky) living in a large outdoor enclosure.

First, we hypothesised social hierarchy to be an important factor in how animals choose their preferred location, with lower ranking individuals’ location being more strongly determined by others’ location than higher ranking ones. Second, we also hypothesised kinships to have an effect, albeit in the opposite direction (Städele, Pines, Swedell, & Vigilant, 2016), resulting in greater predictability of related than unrelated individuals. Third, we hypothesised sex to play a role, such that same-sex individuals determined each other’s position to a stronger degree than opposite-sex individuals, following the idea of male-female role allocation in wolves (Mech, 1999). Lastly, we hypothesised a role of important external events on the stability of the pack. We therefore analysed the predictability of individuals’ locations relative to an upcoming feeding event. Since each individual was chained to a specific location, we predicted that, with increased proximity to feeding time, group stability should decrease.

## Materials and Methods

### Subjects

Our subjects were 15 Siberian huskies (*Canis lupus familiaris*) living as a pack in an open area of 750*m*^2^ (25m × 30m) at the dog-sledding centre ‘Les Attelages de la Roche Percée’ at ‘La Ferme de Nirveau’ in 25510 Pierrefontaine-les-Varans, France (Fig. 1A). Subjects’ ages ranged from 5 to 10 years (M = 6.73, SD = 1.83, Table 1). None of the individuals suffered from any known orthopaedic and/or neurological disorders. Human interactions with the dogs were largely in terms of training for dog sledding and feeding once per day. During the time of recording (October to November 2017) there was no training. Social rank was provided by owners and care-takers based on qualitative assessments (Fig. 1C). Kinship was coded as the presence or absence of siblings in the group (Table 1).

**Table 1:** Identity and social information of individuals.

| **Name** | **Age** | **Rank** | **Sex** | **Siblings** |
| --- | --- | --- | --- | --- |
| Dusty | 10 | 1 | m | Démon, Dubble |
| Démon | 10 | 2 | m | Dusty, Dubble |
| Jack | 5 | 3 | m | Torok, Kinai, Quebec |
| Torok | 5 | 4 | m | Jack, Kinai, Quebec |
| Kinai | 5 | 4 | m | Jack, Torok, Quebec |
| Quebec | 5 | 5 | m | Jack, Torok, Kinai |
| Dubble | 10 | 6 | m | Dusty, Démon |
| Gribouille | 6 | 1 | f | Siska, Babou |
| Babou | 6 | 2 | f | Gribouille, Siska |
| Siska | 6 | 2 | f | Gribouille, Babou |
| Laika | 8 | 3 | f | Elska, Friskies |
| Friskies | 8 | 3 | f | Laika, Elska |
| Tara | 6 | 4 | f | Banquise |
| Banquise | 6 | 5 | f | Tara |
| Elska | 8 | 6 | f | Laika, Friskies |

**Figure 1:**
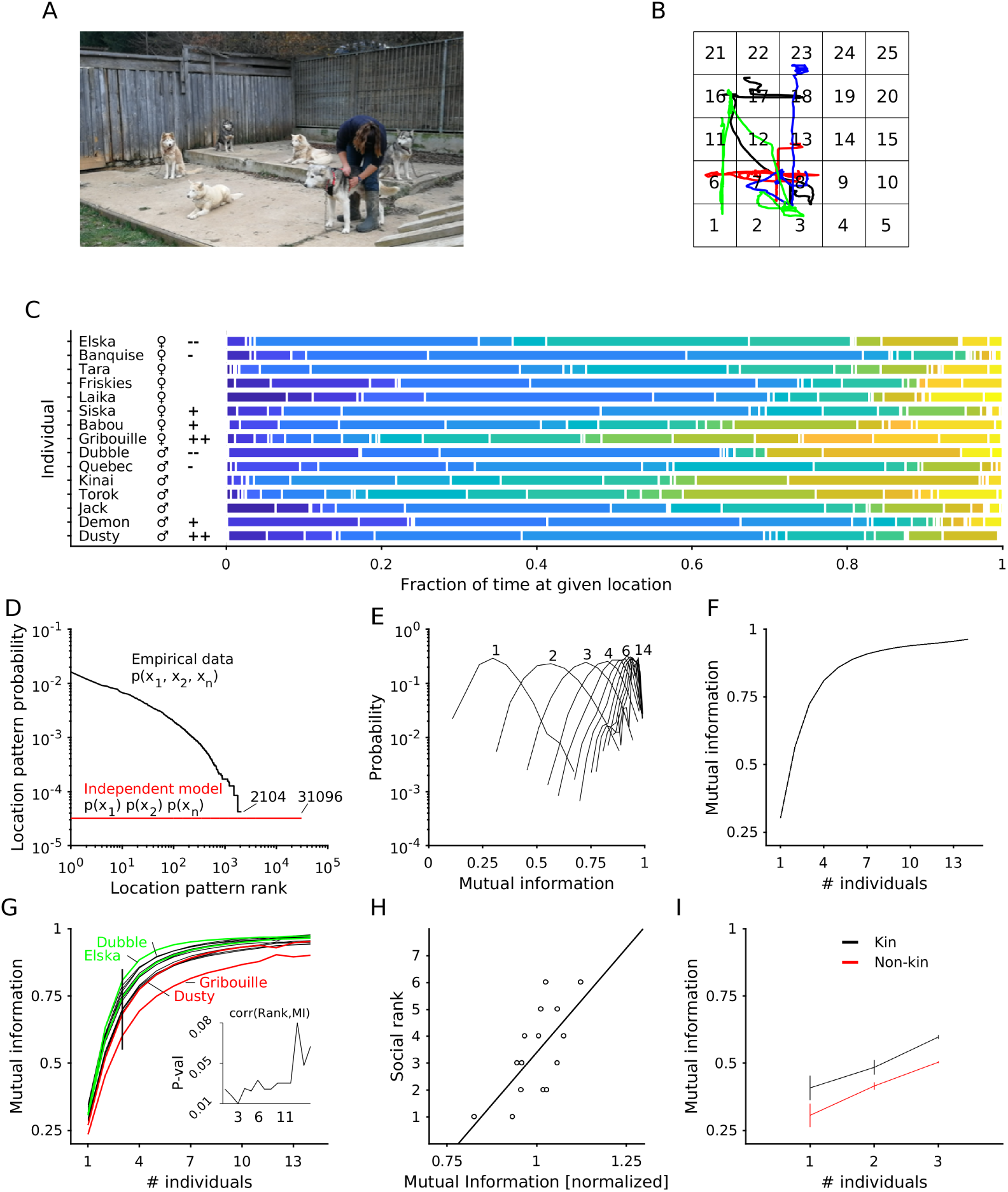
Preparation phase and results of dog tracking. A. Preparation phase: dogs were equipped with GPS loggers mounted to a collar. B. Example movement trajectories of three dogs overlaid onto the 25 patch-grid. C. Fractions of time at given location were plotted for each dog. Colour-coded the locations. Symbols indicate the sex and rank of the individuals. ‘+’ symbols indicate dominance; ‘-’ symbols indicate submission. D. Location patterns were plotted according to the number of occurrences with the most probable location pattern first and the least probable location pattern last (x-axis) and the probability of each location pattern (y-axis). E. The mutual information between the dog *i* (to be explained) and a varying number of other dogs (ranging from 1 to 14) is shown in histograms. F. The median of each histogram (mutual information) in E is plotted against the number of individuals used to explain dog *i*. G. Mutual information is shown as a function of increasing number of individuals used to explain dog *i*. Dominant (red) and submissive (green) individuals, which were subjected to be explained by others, are highlighted. H. The social rank is plotted against and correlated with the normalized mutual information. Normalization was done by dividing the values by the mean. I. Mutual information is plotted against a varying number of kin or non-kin (control). Maximal number of kin is limited to four; hence number of individuals to explain dog *i* is limited to three.

### Data logging

All 15 individuals were fitted with data loggers that provided location data via the global positioning system (GPS), as latitudes and longitudes in degrees, minutes and seconds. As loggers we used ‘Adafruit Ultimate GPS Featherwing’ devices with an update rate of 5Hz, allowing positional accuracy of 1.5 m, a velocity accuracy of .1 m/s and ‘Adafruit Feather Adalogger’ with an ATmega32u4 clocked at 8MHz (Adafruit Industries, NY 10013, USA). Data were locally stored on SanDisk Ultra 16GB MicroSD cards (Western Digital Technologies, Inc., Milpitas, CA 95035, USA). We further used 3.7V 900mAh LiPo batteries to provide power of up to a full day of continuous recording. The hardware was programmed using ‘Arduino Adafruit GPS Library’ to read out the unparsed NMEA sentences. Time stamps were recorded using an in-built real-time clock (RTC).

### Data collection and analysis

We recorded data during three sessions on separate days, lasting nine hours each. In each session we recorded positioning data from all fifteen dogs. We first parsed the NMEA sentences into latitude, longitude and time. We then applied a rotational transformation in the image plane to align the shape of the habitat to an upright rectangle. The values were then converted into a normalized representation of space ranging from 0 to 1 on both axes, while maintaining the spatial proportions. We then equally spaced the habitat into 5 by 5 patches (smaller areas) and converted the normalized GPS locations into the corresponding patch within which the GPS location fell (Fig. 1B).

### Location patterns

For each time point *t* we determined a location pattern consisting of the location patches of all individuals (e.g., location pattern: (x_1_, x_2_, x_3_,… x_n_) = [2, 1, 9,… 4]), where x_i_, is the location of dog *i* (ranging from 1 to 15) and x is the location (ranging from 1 to 25) (Shemesh et al., 2013). We then determined the frequencies of finding the fifteen individuals in given location patterns (p_empirical_ = (x_1_, x_2_, x_3_,… x_n_)) and compared these frequencies with an independent model, assuming that each individual dog moves freely and solely according to its individual preferences (p_independent_ = p(x_1_) p(x_2_) p(x_3_)… p(x_n_)).

### ‘Mutual information’

We based our analyses on the concept of ‘mutual information’ (Shannon, 1948). To this end, we first calculated the uncertainty of the location of dog *i*, defined as the entropy of the distribution of locations of dog *i*, which is

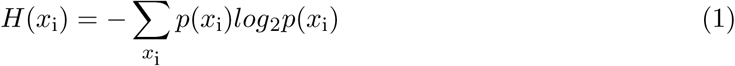

We then calculated the conditional entropy of locations of a pair of dogs (*i, j*) and subtracted that from one of the dogs (*i*); that is, the mutual information between the locations of dog *i* and dog *j*, described as follows:

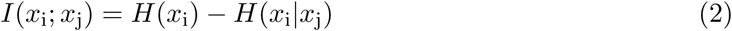

The mutual information for dog *i* and dog *j* was then divided by the entropy of dog *i* to get a normalized measurement of the uncertainty about the location of dog *i*, when considering the joint positions of dogs *i* and *j*. Accordingly, we calculated the mutual information of one dog considering the joint locations of multiple dogs, given by the following equation:

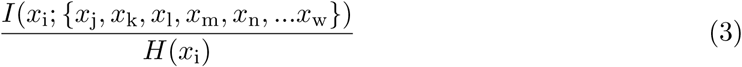

To determine the mutual information between fifteen dogs and 25 possible locations, we used an urn model approach (Sprott, 1978), by selecting 4’000 randomly determined configurations of dog *i* (to-be-explained individual) and dogs *j* to *w* (number of explaining individuals was varied from 1 to 14).

### Predicting individual location: rank and kinship

Mutual information values were binned according to the individual subjects and number of group members taken into consideration to explain a subject’s location. This allowed us to systematically describe dependencies of a subject’s social rank and the predictability of its location by the locations of others. Using Spearman’s rank correlation we hypothesized that low-ranking individuals, more than high-ranking individuals, were subjected to the positions of others and, hence, more predictable regarding their locations. Mutual information values were also used to describe mutual information between siblings and non-relatives. Here, we predicted that mutual information was larger between sibling than non-relatives, given that siblings build a social bond and stay in close proximity to each other. To test this, we performed a two-way analysis of variance with mutual information values as the dependent variable, ‘Kinship’ (yes, no) and ‘# individuals’, describing how many individuals’ locations were used to explain the locations of the individual to be explained, being the grouping (independent) variables.

### Social network

We determined the social network (Fig 2AB) by calculating linkages (edges) between pairs of individuals (nodes). Linkages were defined as mutual information larger than average of all mutual information values. In other words, edges indicated that two individuals (nodes) were influencing each other in terms of their locations above average. The underlying algorithm was a so-called ‘force-directed placement’ plotting routine (Fruchterman & Reingold, 1991). These algorithms follow an aesthetical principle, i.e., they aim at placing all nodes in a way that the edges are close to equal lengths while minimising the number of edge-crossings. Based on the relative positions of sets of nodes and edges, the algorithm assigns forces to them and repositions the sets of nodes and edges to minimize force energies (Kobourov, 2012). In detail, forces push or pull the nodes closer together or further apart, eventually reaching a state of equilibrium, where the spatial repositioning of nodes becomes ineffective, i.e. nodes do not move anymore. Based on these positions, the 2D-graphs were drawn and the sum of edges on the shortest path between individuals with respect to intrinsic attributes, such as sex and kinship, were calculated. We also performed a randomization test by computing the sum of edges of randomly assigned individuals to the current positions, shuffling kinship labels while preserving the existing node locations. We ran this iterative procedure 200 times and compared the actual values with it. We computed a similar procedure for sex, reassigning male and female labels to current positions, in order to calculate same- and opposite-sex distances under randomly assigned conditions.

**Figure 2:**
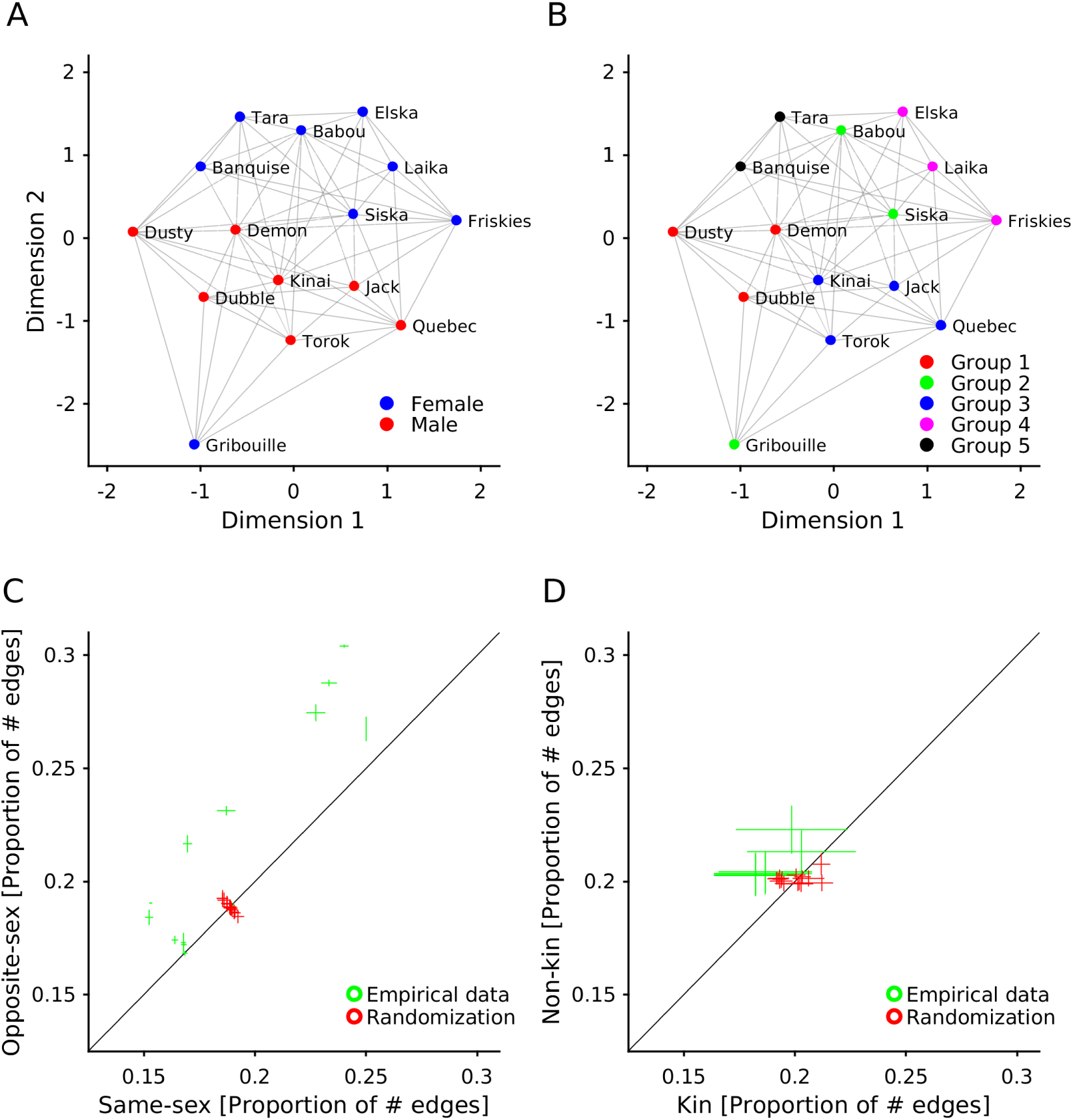
Social networks. A. shows a 2D map illustrating the social network of mutual influence between members. An increasing number of links indicate an increased influence regarding dog *i* ‘s position through others. Colour-coded is the sex of the individuals. B. Same as in A. Colour-coded is the kinship, indicating the siblings. C. The sum of edges on the shortest path between individuals of same and opposite sexes, as extracted from A, are shown. Means and standard deviations are indicated by green lines. Red lines denote the results from the randomization procedure. Numbers mark the number of individuals that were used for the mutual information computation. D. Same as in C, but according to kinship.

### External event

The most relevant non-social event in subjects’ daily activities was most certainly the advent of feeding, which always took place around 18:00 local time. For this purpose, the dogs were individually tethered to a chain spot on the ground in order to avoid food-related aggression. Therefore, recording sessions regularly ended around this time, when the loggers were turned off and the harnesses taken off over night. The actual feeding event and its influence on group structure, for this reason, could not be investigated directly, although we could study the effects of anticipation by time-logging the entire recording session towards the feeding event. We subdivided the entire session into time frames of 54 min and calculated group information within these time periods. We applied Gaussian Kernel linear regression on time frames (elapsed time) and mutual information values.

## Results

Preferences for locations were measured for each individual as fractions of time (Fig. 1C). Probabilities of location patterns of the empirical data and of the independent model were highly different (Fig 1D). Of possible 25^15^ states (i.e., 25 locations by 15 individuals), only 2’104 states (i.e. location patterns) were observed. However, the observed states occurred at high frequencies, up to 382 times (Fig 1D, black line). In contrast, the individual model predicted roughly 31’096 location patterns, however, at lower probabilities (i.e. each location pattern occurred one time only) (Fig 1D, red line). Hence, our data indicate that the group restricts the number of socially accepted location patterns to about 7% for the average individual (2’104 out of 31’096 location patterns). The remaining 28’992 location patterns, predicted by a model based on individual preferences, were socially avoided.

We then quantified the strength of dependencies between pairs of individuals. In detail, we addressed the extent to which knowing the location of one dog helped inferring the location of another dog. In a scenario where two individuals were fully independent on each other, knowing one individual’s location would not make any predictions about another individual’s location. We therefore determined the entropy (i.e. uncertainty) of locations for each individual to predict the current location based on previous location information, and the mutual information between two individuals, i.e. how much of the uncertainty about the location of one individual could be reduced by knowing the location of another individual (see methods). We also calculated the joint locations of multiple individuals to explain the locations of single individuals. The results showed that knowing the locations of one individual allowed predicting a second individual’s location in a range from 11 to 67%, with a median of 30.2% (see Figure 1E). Knowing the location of two individuals allowed predicting the location of a third individual for up to 56.5% (median). With increasing number of individuals’ joint locations, the explanatory power of a further individual increased, reaching a median of 96.3% with 14 individuals (Figure 1F).

### Social rank

We then sorted the mutual information values according to the individual explained (e.g. dog *i*), independent of the number of individuals used to explain that individual (e.g. joint information of dog *j*, dog *k* and dog *l*). We found that two individuals (Dubble and Elska), determined as submissive by the owners, turned out to be strongly influenced by other dogs’ locations in terms of choosing their own locations (Figure 1G). Two further individuals (Dusty and Gribouille), classified as the most dominant dogs in the pack, were found to be least influenced by other dogs’ locations in terms of choosing their own locations. We correlated the social rank and the mutual information values and found a positive correlation for most group sizes (Figure 1G, inset). The black line in Figure 1G indicates that group size of four individuals (i.e. one individual to be explained and three individuals to explain) showed the strongest correlation between social rank and mutual information values (rs(13) = .64, *p* < .05) (Figure 1H).

### Kinship

We also calculated the mutual information for siblings. Here, we determined the degree to which we could explain an individual’s location by the location information of all its siblings. As a control, we calculated the predictability of the same individual but with non-kin individuals contributing to the joint location information. We found that individual location could be predicted to a larger degree by the location of siblings than unrelated group members (Kinship: F(1,1446) = 7.88, *p* < .01, Mean sq = .24; # individuals: F(2,1446) = 127.82, *p* < .001, Mean sq = 3.89) (Figure 1I).

### Social network

We calculated the linkages between individuals based on the definition that a linkage exists if it is above average. Social networks were determined using the ‘Force’ plotting routine (Fruchterman & Reingold, 1991), in our case based on mutual information values derived from four individuals (i.e. one individual to be explained and three individuals to explain, Figure 2AB). Results showed that nodes and linkages grouped according to the sex of the individuals (Fig 2A), with sums of edges on the shortest path between individuals of same and opposite sexes differing significantly (F(1,392) = 1.93, *p* < .05, Mean sq = 0.007). The number of individuals used to explain a single individual turned out to be a modulating factor (F(13,392) = 1.83, *p* < .05, Mean sq = 0.006) (Fig 2C). In an iterative t-test comparison (corrected for multiple comparisons), we found that same-sex individuals were closer to each other than predicted by a randomization control procedure when the social network was constructed on the basis of mutual information values computed by the locations of 3 to 5 individuals to explain the locations of another single individual (all *p* < .01). Similarly, the same social network revealed a grouping according to kinship, where siblings were spatially closer than non-siblings (F(1,392) = 1.77, *p* < .05, Mean sq = 0.008) (Fig 2BD). The number of individuals required to explain a single individual’s location was a modulating factor (F(13,392) = 2.31, *p* < .01, Mean sq = 0.007) (Fig 2D). Critically, siblings were closer than predicted by an iterative procedure randomizing kinship relationships when the social network was constructed on the basis of mutual information values computed by the locations of 3 or 6 and more individuals to explain the locations of another single individual (all *p* < .05). The individuals Gribouille and, to a smaller extent, Dusty were positioned slightly off from the general cluster. These individuals are dominant to all others and have fewer dependencies on other individuals. Further Gribouille is primarily linked to male individuals (Figure 2A) and appears to be separated from her own kin (Figure 2B).

### Anticipating non-social events

Lastly, we determined the extent to which the group dependency was affected by external events, introducing an interference factor on the group formation. We therefore subdivided the recording session according to the relative time prior to feeding, which regularly occurred around 18:00 local time, and determined the mutual information between each individual with each other individual and each other pair or group of individuals, as described above, with regard to these particular time windows. We found that the closer the time progressed toward feeding, the smaller the probabilities of predicting one individual’s position by looking at other individuals’ positions became (Figure 3A). We interpret this as a loss of group stability due to the forthcoming feeding event that led to a group structure with fewer social constraints. The greatest mutual information values, hence the strongest dependencies, were found during a mid-day period. In contrast, early morning and evening periods – the latter being close to feeding – showed less strong dependencies, hence reduced stability of the pack (Fig 3B). Interestingly, this effect was strongest for mutual information values determined from pairs of animals (i.e. addressing the question “how much of location of dog *i* can be explained by location of dog *j* ?”). In contrast, mutual information values are more constant throughout the day when determined based on a large group of animals (e.g. how much of the locations of dog *i* can we explain by looking at the remaining 14 dogs?).

**Figure 3:**
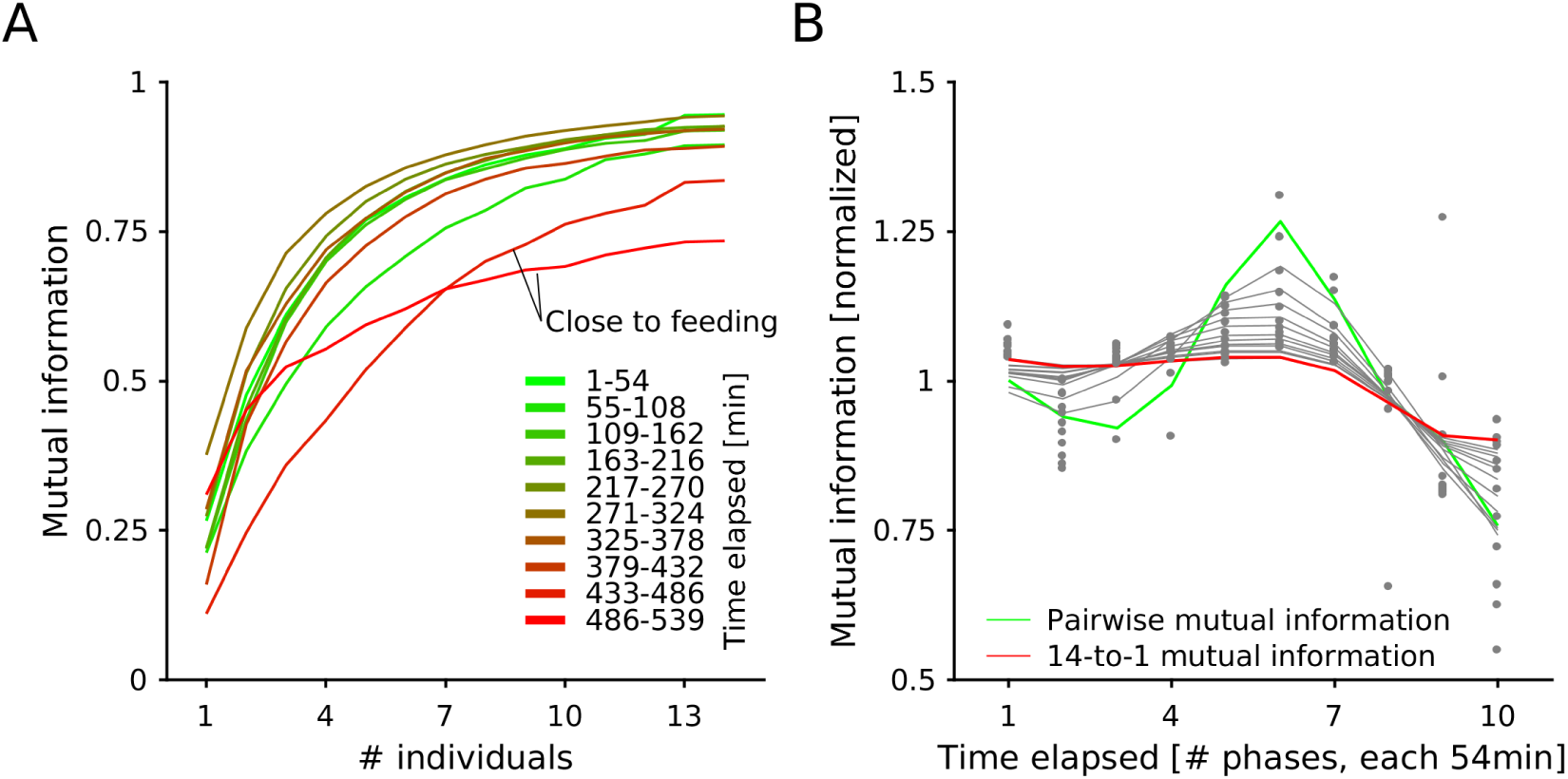
Influence of external event. A. Mutual information values extracted from various time periods are plotted against the number of individuals used to explain dog *i*. The colour-code indicates the elapsed time from the start of the recording. ‘green’ time segments are closer to the onset of recording, while ‘red’ time segments are close to an upcoming feeding event and therefore closer to the end of the recording. B. Normalized mutual information is plotted against the elapsed time from onset of recording. Normalization was done by dividing the mutual information values by their mean. We applied a Gaussian Kernel linear regression function with a Kernel bandwidth of 1 to describe the data.

## Discussion

We described movement patterns in a pack of dogs by collecting GPS (NMEA) data from multiple animals in parallel and, using a real time clock (RTC) on each logger, synchronising the individual movement patterns. We relied on a classic mathematical theory of communication (Shannon, 1948) to quantify the extent to which one individual’s movements can be explained by other individuals’ movements. We found highly correlated group behaviour, first, insofar as the number of observed location patterns was very restricted and by a multitude smaller than what would be predicted if individuals freely moved according to their individual preferences. Hence, ‘socially allowed’ configurations are governed by the group, limiting individual freedom in movement to about of 4%, suggesting that individuals’ movement decisions were determined by others up to 96%.

We found that dominant individuals were influenced by other individuals to a lesser degree than subordinate ones. We also found that, while same-sex individuals influenced each other more strongly than opposite-sex individuals, the pack was spatially controlled by two dominant individuals; an alpha male and an alpha female. Furthermore, we found that kin relationships increased the mutual dependencies of individuals, with siblings moving together, a social structure similar to family composition in free-ranging wolves (Gadbois, 2004; Mech, 1999; Packard, 2003). Interestingly, the level at which grouping effects emerged varies depending on the attributes: Social rank seems to play a major role in very small configurations of animals, emerging already in the mutual information between one individual and two other individuals. Sex classes, too, emerged at early level of co-dependencies, i.e. with groups of 4 to 6 animals. In contrast, kinship classes are more strongly subjected to the organization of the whole pack, emerging most prominently at 7 and more individuals.

We also showed that group structure responded to an external factor, that is, upcoming feeding events. Interestingly, as time progressed the mutual information between animals decreased, suggesting reduced network stability due to the upcoming feeding event. This is to some extent surprising in that it does not reflect food competition by means of dominating others, but disintegration of social structure. Alternatively, the findings might be interpreted by considering the given feeding ecology, where food is available on a daily basis and guaranteed to each individual, irrespective of social rank and sex.

### Evolution of social structure

Studies in wild and captive wolves disagree on what should be considered the natural social and hierarchical structure. While captive wolves and domestic dogs showed heightened agonistic behaviour and a social structure best described by a linear hierarchical model (Cafazzo, Lazzaroni, & Marshall-Pescini, 2016; Cafazzo, Valsecchi, Bonanni, & Natoli, 2010), wild wolves have been described as living in family units, consisting of a breeding pair with their offspring (Gadbois, 2004; Mech, 1999; Packard, 2003). Family compositions suggest a non-linear and more complex but, at the same time, more flexible hierarchical structure (Packard, 2003) with the parents playing the roles of leaders, making group decisions and initiating group movements (Peterson, Jacobs, Drummer, Mech, & Smith, 2002). Little is known about social behaviour, dominance and rank in natural packs of dogs. Free-ranging dogs normally build packs ranging from 2 to 8 individuals, in rare cases over 20 individuals (Bonanni & Cafazzo, 2014), and consist of males and females. Females tend to have multi-male mating preferences (Cafazzo, Bonanni, Valsecchi, & Natoli, 2014) and subsequently raise their offspring. Also noteworthy is that mating strategies, raising preferences and rank hierarchies vary in dogs and wolves depending on the feeding ecology (Marshall-Pescini, Cafazzo, Viranyi, & Range, 2017).

Importantly, in our study, we used human-socialised dogs, as opposed to stray dogs with unknown socialisation histories. Subjects had been raised in human social environments, but kept in a relatively free-ranging and natural pack. As a consequence, their social structure might differ from both wolves and free-ranging stray dogs, given the absence of influencing factors such as foraging/hunting, mating/breeding and predators/danger sources, and the presence of factors such as human-dependency, controlled feeding and care taking.

### Conclusions

With recent developments in hard- and software technologies, traditional methods in animal behaviour research are likely to experience fundamental changes towards automated continuous recording procedures, allowing the quantification of not only single-animal activities, but interactions of groups of animals with an unprecedented temporal accuracy. While many algorithms to quantifying group interactions are highly sophisticated and computationally intense (Bialek et al., 2012; Lukeman et al., 2010; Vicsek et al., 1995), we here present a relatively simple mathematical approach, rooted in information theory (Shannon, 1948) devices. We were able to describe the social dynamics in a semi-natural pack of freely interacting dogs to a surprisingly complex degree, which revealed the influence of evolutionarily inherited social structures that included kin relations and dominance relations, as well as predictions made by individuals about relevant forthcoming external events.

## Acknowledgements

We thank ‘Les Attelages de la Roche Percée’ at ‘La Ferme de Nirveau’ in Pierrefontaine-les-Varans, 25510, France, for their support. We are grateful to the Swiss National Science Foundation for supporting this project via the Ambizione Fellowship (PZ00P3 154741) awarded to CDD. KZ has also been supported by the Swiss National Science Foundation (31003A 166458). We thank Guillaume Dezecache and Malte J. Rasch for comments on the manuscript.

## Author contributions

CDD: study design, data collection, analysis and interpretation, writing article, provision of necessary tools; EF: data collection, provision of resources; KZ: provision of necessary tools and resources, writing article.

## Ethical approval

According to the local authorities (Comité d’Ethique de l’Expérimentation Animale Grand Campus Dijon, Université de Bourgogne, Maison de l’Université, Esplanade Erasme, 21078 Dijon, France), non-invasive studies on dogs are allowed to be conducted without any special permission in France. ‘Les Attelages de la Roche Percée’ at ‘La Ferme de Nirveau’ in 25510 Pierrefontaine-les-Varans, France, responded to our enquiry and volunteered to participate in this study.

